# Thermodynamic and Kinetic Characterization of Protein Conformational Dynamics within a Riemannian Diffusion Formalism

**DOI:** 10.1101/707711

**Authors:** Curtis Goolsby, Ashkan Fakharzadeh, Mahmoud Moradi

**Affiliations:** Department of Chemistry and Biochemistry, University of Arkansas, Fayetteville, AR 72701, U.S.A; Department of Physics, North Carolina State University, Raleigh, NC 27607, U.S.A

## Abstract

We have formulated a Riemannian framework for describing the geometry of collective variable spaces of biomolecules within the context of collective variable based molecular dynamics simulations. The formalism provides a theoretical framework to develop enhanced sampling techniques, path-finding algorithms, and transition rate estimators consistent with a Riemannian treatment of the collective variable space, where the quantities of interest such as the potential of the mean force, minimum free energy path, the diffusion constant, and the transition rate remain invariant under coordinate transformation due to the Riemannian treatment of the collective variable space. Specific algorithms within this framework are discussed such as the Riemannian umbrella sampling, the Riemannian string method, and a Riemannian-Bayesian estimator of free energy and diffusion constant, which can be used to estimate the transition rate along a minimum free energy path.

## I. INTRODUCTION

Biomolecular simulations have made enormous progress in recent years. The advent of Molecular dynamics (MD) in order to study biomolecular phenomena [29, 41, 45] has given researchers new insights into previously un-viewable phenomena. MD has overcome [40, 42] the common limitation of experimental techniques which force the user to choose between high-resolution static or low-resolution dynamic pictures of biomolecular systems.

MD does however have a few drawbacks. First, there is a temporal issue in that MD simulations have a shorter time scale than many relevant biological phenomena. In addition, MD has difficulty producing a full description of high-dimensional free energy landscapes. Various enhanced sampling techniques have been developed over the last few decades in order to overcome these limitations [1, 37, 43, 51, 53, 58, 59, 65, 66, 68, 71]. These techniques are often successful for simple toy models (e.g., dialanine peptide [10, 24, 38, 43, 48, 54]); however, practical applications remain quite challenging.

Perhaps the most obvious line of attack for solving these problems is simply to have stronger computing power and more efficient or specialized computer hardware. In the past 40 years, extraordinary advances have been made in this regard. Current state of the art architectures such as Anton [60, 61], peta-scale supercomputers [2, 14, 15], and GPUs [28, 30, 64, 67] have capabilities that allow for simulations with billions of particles over thousands of processors. The size of supercomputers have also given rise to algorithms developed for enhancing the sampling by brute force (rather than statistical mechanics based enhanced sampling) such as particle mesh ewald [5] and dynamic load balancing [22]. The focus of this article, however, is the enhanced sampling techniques that are specifically based on enhancing the sampling within a statistical mechanical framework.

In order to obtain relevant thermodynamic and kinetic [33] properties of a statistical mechanical system, one must integrate over high dimensional spaces, which requires large sets of independent and identically distributed samples. In order to manage the dimensionality of systems the assumption is often made that the vast majority of the high-dimensional space is practically empty (or occupied by microstates of negligible probability) and the occupied space can be approximated by a lower-dimensional manifold which contains all relevant conformations from stable states to important transition states. Ideally, any elementary reaction, can be thought of as a transition between two stable states, characterized by a transition pathway [11, 23] (or committor function [16]).

Some methods use the ideas of an intrinsic manifold without specifically using a conventional dimensionality reduction technique as in path-optimization techniques [8, 9, 13, 46, 54, 58], which all rely on a localized transition tube. Other methods attempt to identify the intrinsic manifold by using statistical learning methods such as principal component analysis [3], isomap [11], and diffusion map.[23, 25]. Often these techniques are used to analyze MD trajectories [3, 11, 23, 57] or are combined with enhanced sampling as in metadynamics [63, 69] or adaptive biasing force [31].

The use of a coarse/reduced variable space (whether one- or multi-dimensional) in both free energy calculation methods and path-finding algorithms is quite common and various algorithms have been developed to estimate free energies [4, 24, 32, 34, 36, 37, 43, 47, 56, 59, 68] or find transition pathways [8, 9, 13, 46, 49, 54, 58] in such spaces. What is somewhat missing is a theoretical framework that allows for a rigorous treatment of the issues one needs to deal with when working with such methods. For instance, the collective variables used in collective-variable based enhanced sampling methods are often non-linear transformations of atomic coordinates. This complicates their application as noted previously by Johnson and Hummer [39], who show the conventional minimum free energy path (MFEP) obtained from various path-finding algorithms are collective-variable specific and are not invariant under non-linear coordinate transformations. Such difficulties have often been ignored in the past in the majority of the applications of the collective-variable based simulations; however, there has been attempts in addressing them as in the aforementioned work [39] or within the framework of Transition Path Theory [18, 46]. We recently introduced a Riemannian framework for the rigorous treatment of the collective-variable based spaces within the context of enhanced sampling and path-finding algorithms [21], where the distance, free energy, and MFEP are redefined in a way that are all invariant quantities and do not change under coordinate transformations. Our previous work focused on the thermodynamic characterization of protein dynamics within a Riemannian diffusion model [21]. Here we will extend the formalism to kinetics and rate calculation techniques within the same Riemannian framework.

## II. FROM EUCLIDEAN TO RIEMANNIAN DIFFUSION

Euclidean geometry was developed by the Greek Euclid cerca 300 BC. Riemann’s work on generalizing the differential surfaces of IR^3^ led to progress in many fields of science. Further progress upon his ideas allowed for the formulation of Einstein’s General Theory of Relativity [19], progress in group theory [50] and generated differentiable topology.

Riemannian geometry provides a robust mathematical framework to develop a formalism for the geometry and dynamics of collective/coarse variable spaces, defined to reduce the dimensionality of the atmoic models of macromolecular systems. For instance, consider a transmembrane protein whose transmembrane helices rotate under certain conditions to allow oepning or closing of a gate and transporting materials across the membrane. A reasonable collective variable for such a system would be the orientation of the transmembrane helices that can be determined using principal axes of the helices or their orientation quaternions [26, 52]. In order to work in these spaces, we must first have a formalism that allows us to answer common questions which are posed in a typical collective-variable based simulation. Examples of such questions include: what is the distance between two points in the collective variable space? How is the potential of the mean force (PMF) defined at a given point in the collective variable space? How does a biasing potential affect the distribution? How can one find the MFEP? How can we estimate the PMF from biased simulations along the MFEP? What is the diffusion constant along a transition pathway? How can we estimate the rates of a reaction from the PMF and diffusion constants? The questions have previously been answered within a Euclidean framework but with our Riemannian treatment of the collective variable space, some of these concepts and quantities need to be revisited. We will begin to answer these questions one at a time.

Imagine a system containing *N* atoms which are represented based upon their coordinates in three dimensional space, ***x***, their momenta ***p***, and a potential energy surface *V* (***x***). Given that they lie in a canonical equilibrium distribution their motion can be described with 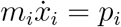 and Langevin Dynamics [44] written here as:

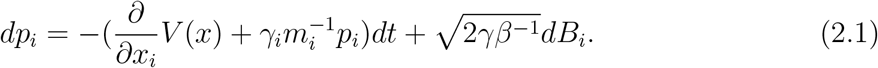

Given the usual assumption for brownian motion, i.e. that the system is overdamped and thus the left side of the equation is zero, we can write:

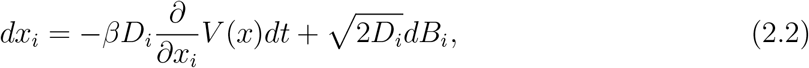

for which *dB*_*i*_ is a Weiner Process and 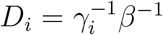 is the diffusion constant for atom i. In order to reduce the variable space to a coarser space we make the relation 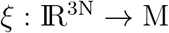 M such that *ξ* is a multi-dimensional variable which reduces the system such that (*M, **g***) is a complete Riemannian Manifold of dimension n with a position dependent metric ***g***(*x*). Thus we can describe an effective potential as:

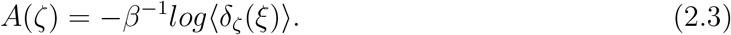

in which *ζ* is any point on M and *A*(*ζ*) is the Riemannian effective potential which is related to Euclidean PMF, *Ã*(*ζ*), by:

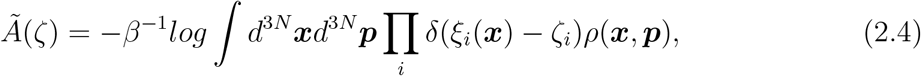

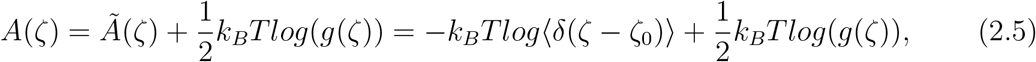

and *δ* is the dirac delta function which in Euclidean and then Riemannian formalisms satisfy (6) and (7).

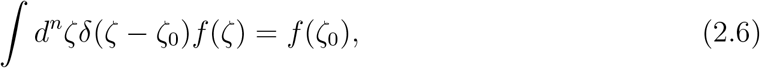

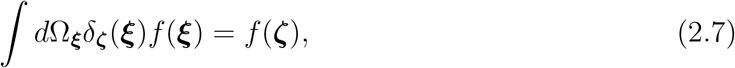

where Ω is the volume element given by 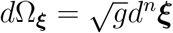. Noticing that both the Euclidean formalism and Riemannian formalism share the same ***ξ*** we can now note that their PMF, is the same. We rewrite (3) as:

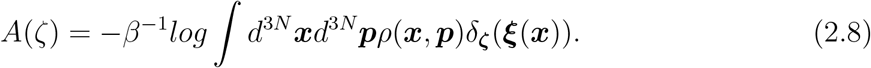

From this we can write, in a similar fashion to (2):

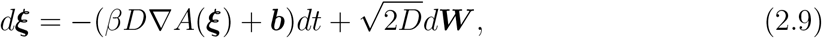

where we have removed the position dependence of the diffusion constant, which may now be represented by an arbitrary constant. Instead the position dependence is coupled into the metric, g, as seen below, and *dW* is once again a Weiner process. In order to continue our formulation we make the assumption that our model is diffusive. This allows us to write our model in terms of the Fokker-Planck (or Smulchowsky) equation [20]:

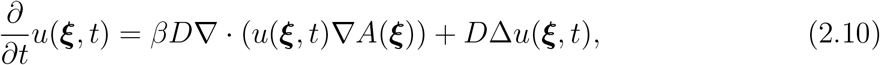

which contains a gradient defined by 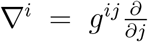, the Laplace-Beltrami operator, 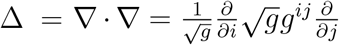, and *u*(*ξ,t*)is the probability of finding the system at *ξ* at time *t* with a boundary condition of *u*(*ξ*, 0) = *u*_0_(*ξ*).

This summarizes our diffusion model, which was previously introduced in Ref. [21]. In the following we derive a new relation that allows the estimation of the PMF, metric, diffusion constant, and transition rate from unbiased simulations performed along an approximate MFEP.

## III. TRANSITION RATE ESTIMATION

We will later discuss how one can find a MFEP using path-finding algorithms. For now, let us assume we have found such a pathway and we have placed a system on a given point close to this pathway at time *t*. One can write:

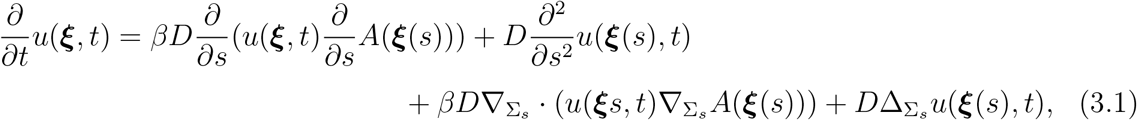

where the variable *s* is the geodesic distance along the MFEP ***ξ***(*s*), where it parametrizes ***ξ***(*s*). ***κ*** is defined on the *n* − 1 dimensional subspace of ***ξ*** perpendicular to ***ξ***(*s*) (with its origin placed on ***ξ***(*s*)). 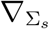 denotes the ∇ operator in the subspace of ***κ***. We have thus projected ***ξ***(*s*) onto two components including the one-dimensional *s* and the *n −* 1 dimensional ***κ***.

Let us now define 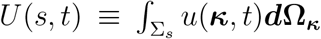, where Σ_*s*_ is an arbotrary portion of the subspace of ***κ*** around its origin. On the MFEP, let us also define a univariate PMF *G*(*s*) = *A*(*ξ*(*s*)). We note that on a converged pathway, 〈*κ*〉 = 0, gives a zero drift pathway in any high dimensional space.

Continuing, we can integrate over individual terms in Relation (3.1), where the LHS of the equation becomes 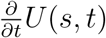 and the RHS terms can be approximated as follows. Assuming *u*(***ξ**, t*) is much larger within a relatively narrow “tube” around the MFEP, we can integrate over the cross section of this tube and the areas around the tube. In other words, we choose Σ_*s*_ to be a portion of the ***κ*** space that covers the cross section of transition tube that falls within this space as well as some low-probability areas around it. The first term of the RHS of Relation (3.1) will thus be:

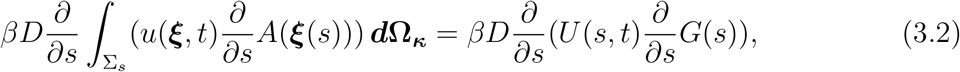

where we define a univariate PMF *G*(*s*) = *A*(*ξ*(*s*)) and assume *A*(*ξ*(*s*)) is more or less constant on any cross section of transition tube perpendicular to the MFEP. The next term becomes 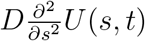 after integration and the third term will vanish assuming 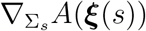 stays close to zero within the tube. To be more precise, we assume the following integral is negligible as 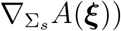 is negligible close the MFEP:

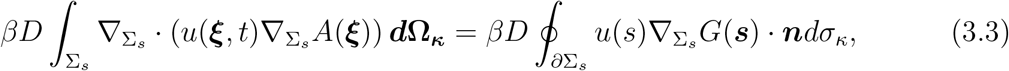

where the divergence theorem is used. Similarly, we can use the divergence theorem to reduce the last term to:

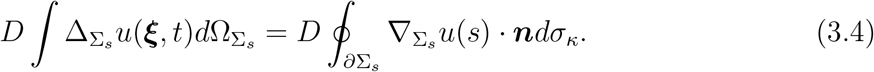

We assume this term is also negligible since the 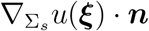 term is only evaluated outside the transition tube and is expected to be close to zero on average. These approximations reduce Relation (3.1) to a one-dimensional diffusion equation in terms of probability density *U* (*s, t*) and the univariate PMF (or potential energy) *G*(*s*):

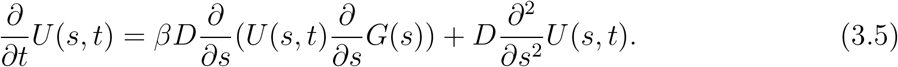

The thin transition tube, used to simplify Relation (3.1) is a common assumption made for path-finding algorithms such as string method [17]. At this point, we have reduced our atomic model, first to a coarse variable space, and then we have focused upon a single dimension, namely the transition pathway of interest.

Relation (3.5) is quite similar to the conventional Smoluchovsky equation in a one-dimensional Euclidean space. However, this is due to the fact that we assumed *s* to represent the geodesic distance along the MFEP path. A slightly more general relation can be derived based on (3.5) for an arbitrary parameter *r*, parametrizing the MFEP path. The *r* space will then be associated with a 1D metric *h*(*r*). We have *ds*^2^ = *h*(*r*)*dr*^2^ or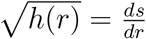. The Riemannian Smoluchowsky equation in then locally written in terms of *r* as:

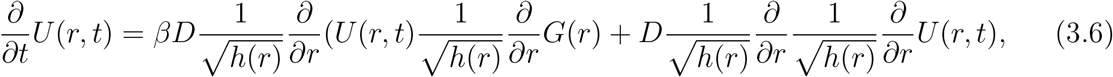

which is slighhtly different from a conventional 1D position-dependent diffusion equation. Here *D/h*(*r*) is equivalent to the conventional 1D position-dependent diffusion constant and *G*(*r*) is the same as the conventional 1D PMF in terms of *r* with an extra term 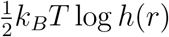, which makes the Riemannian PMF independent of the choice of *r*.

Finally, we can rewrite Relation (3.6) as:

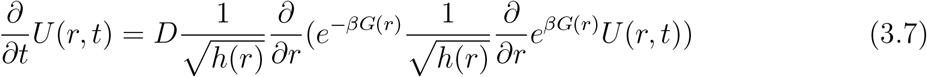

Now suppose that one has been able to identify the MFEP without necessarily quantifying the metric. Relation (3.7) provides a framework to determine the dynamics of the system as long as it stays close to the MFEP transion tube. The PMF and metric along *r* (*G*(*r*) and *h*(*r*)) fully describe the diffusive dynamics and the arbitrary diffusion constant *D* simply determines the unit of *h*(*r*). In Section V, we will discuss in more detail how the estimation of *G*(*r*) and/or *h*(*r*) is possible using unbiased or biased simulations; however, none of these discussions would be relevant without identifying the MFEP.

## IV. PATH FINDING ALGORITHMS

In Sections II and III, we developed the Riemannian framework of path finding algorithms. The PMF defined above is invariant under coordinate transformation as is the MFEP. The MFEP and the formalism of the String Method with Swarms of Trajectories, SMwST [55]. We will also in this section describe Post-Hoc String Method which allows a user to define certain parameters to move from state a to state b while finding a minimum free energy pathway, MFEP [27, 46], by defining a string based on previously performed sampling.

First we present PHSM. In order to explore the reaction coordinates we need to find a MFEP. We can find the MFEP by using a predetermined sample set to obtain a reaction pathway. Before using PHSM the user must supply a large set of conformations and weights for said conformations. PHSM then requires inputs which give (i) a metric/distance norm ||.|| to measure the closeness of conformations; (ii) an initial pathway of n conformations; (iii) transition tube thickness parameter *ϵ*; and (iv) convergence criteria including a threshold *δ* and a persistence number P: The particular algorithms used below for finding an initial path and re-parametrization have their own parameters (i.e., ∆min/max and *δ′*, respectively). In order to completely clarify this process, we will describe the entire process now.

The collective variable space and metric are defined as above. Next convergence criteria must be selected, ∆min/max, *δ*, *δ*’, and *ϵ*. The ∆min/max determine the spacing of your images. The drift, which can be alternatively defined depending on your variable space must be less than *δ′*, and finally the transition tube has a thickness of *ϵ* and must move from its original center by no more than *δ* which has a usage we will see in a moment. The ∆min/max are then used to generate an initial string which varies from *ζ*(0) to *ζ*(1) over the coarse variable space with distance between images defined by ∆min/max. We note that the values chosen are dependent on the distance metric used. This serves as a seed for the string method that follows. This initial string and subsequent strings are updated with new centers found by Voronoi Tesselation [70]. These centers are rejected if they move more than the *ϵ* or *δ* parameters, i.e. if the center strays from the initial center by more than the tube thickness or if the value strays too much from the previous center the value will be discarded. If all the centers are discarded, parameters must be adjusted. The re-parameterization of image centers is performed with de Casteljaus algorithm [12] in order to implement Bezier curves [6, 7] for interpolation. Finally, if the change in the string centers from the previous step is less than *δ′*, the string is converged.

Now that we have a seed string from PHSM, we will discuss SMwST. SMwST starts from an initial string, defined by N points (or images/windows) denoted *x*_*I*_, I=1, …, N. A collective variable *ξ* is defined as above and the initial image centers are given by the initial string. A swarm of trajectories are created, giving M copies of each point or N x M systems in total. Each SMwST system then undergoes three iterative steps. First, the system is restrained for a defined number of steps, *τ*_*r*_, using a harmonic potential, 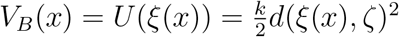, which has its center on the current center and the distance metric defined as *d*(*x, x*_0_). The simulation is then released from their harmonic potential and allowed to drift for a defined number of steps, *τ*_*d*_. Finally, the system is re-parameterized to a new center based upon the average *ξ*(*x*(*τ*_*r*_ + *τ*_*d*_)) values for all M systems associated with an image. Images are kept equidistant and new systems are created until the method reaches convergence to a zero-drift path, i.e. the string centers oscillate. In contrast to a Euclidean SMwST which does not necessarily identify the zero-drift path in the normal coordinates our Riemannian SMwST identifies the zero-drift path in the normal coordinates which provides a coordinate transformation invariant solution.

Our version of SMwST varies in four distinct points from the conventional SMwST:

1. The metric at each image center *ξ*_*I*_ is estimated via:

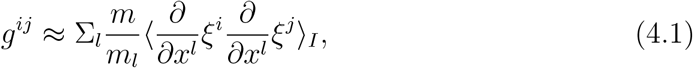

in which the average over the images is calculated for each iteration by using the *x*(*τ*_*r*_) values observed for all M copies of I.
2. The biasing potential is approximated by using *g*(*ξ*_*I*_) to calculate the inner product of 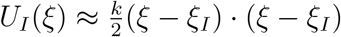.
3. As opposed to using Euclidean distance to perform the interpolation necessary to update images, the distance between two consecutive images is defined by 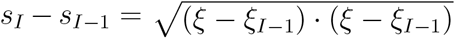, in which *g*(*ξ*_*I*−1_) is used to evaluate the inner product.
4. In the re-parametrization step, the mean drifted centers 〈*ξ*(*τ*_*D*_)〉, used in conventional SMwST, are shifted by a biasing term *−**b***(*ξ*_*I*_)*τ*_*d*_∆*t* for each image I. 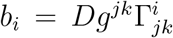 is used to evaluate the geometric drift, where the Christoffel symbols are defined as 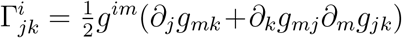. The metric derivatives can be approximated by finite difference, e.g., 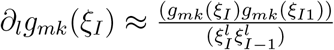.

## V. PERTURBED FREE ENERGY VS PMF

As in the conventional case, the string centers can then be used as the harmonic centers for US simulations. From this the US trajectories can be used to estimate the perturbed free energies by self-consistently solving:

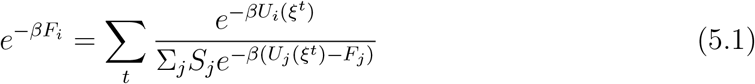

For which the sums are over all coarse variables and the images. F is an estimator for perturbed free energy and 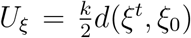 are the harmonic potentials calculated, around the point *ξ*_0_ with a distance metric *d*(*ξ, ξ*_0_), as described in the following five equations.

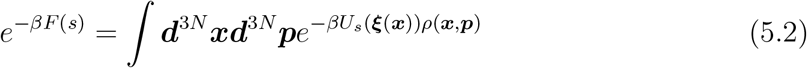

Or

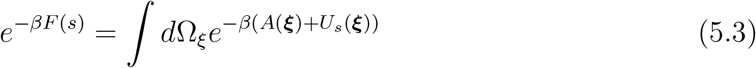

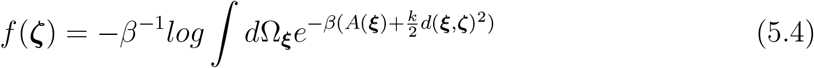

Unlike *F* (*s*), which is defined on the *ζ*(*s*) curve only, *f* (*ξ*) is defined everywhere on the manifold; however, the two functions have the same values at any point along the curve, i.e., *f* (*ζ*(*s*)) = *F* (*s*). Following a similar approach as in the 1-D case we can employ a Riemannian adaptation of the Weirstrauss transform to derive a meaningful relation between the Riemannian PMF and the perturbed free energy.

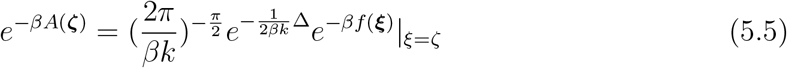

Given a large force constant, k, equation 24 can be simplified to

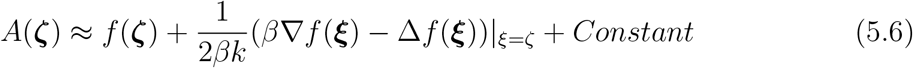

## VI. DETERMINING THE RATE OF TRANSITION

Our final formulation mathematically follows closely the work of Hummer [35], except we are working within a Riemannian geometry. We define 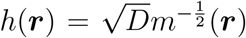 and *H*(***r***, *t*) = *e*^*βG*(***r***)^*U* (***r**, t*) such that (14) can be rewritten in an easily discretized form:

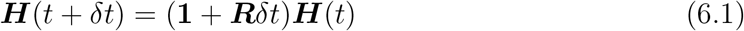

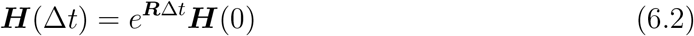

***H*** is here a discretized representation of the transition probabilities which can be determined empirically and R is a tridiagonal matrix for which more can be seen in section 4.

We can self consistently [62] solve by maximizing the likelihood between states at t and t*±*1 defined by:

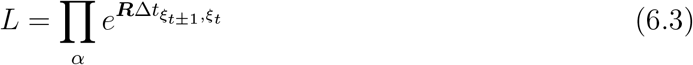

using the tridiagonal ***R*** from (16):

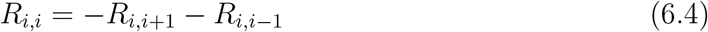

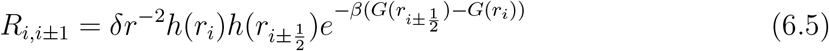

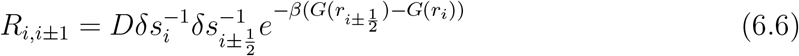

using the relation that 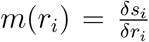 and r values chosen such that *δr*_*i*_ = *δr*_*i*+1_. Now it can be seen that working in a Riemannian collective variable space is valid for the determination of quantities of interest such as rate, free energy calculations, diffusion, and reaction pathways.

## VII. ORIENTATION QUATERNION

Among the different routes to describe rotation and translation, perhaps using quaternions are more convenient one. In our approach, quaternions are 4-vector (*q*_0_, *q*_1_, *q*_2_, *q*_3_) which are normalized, 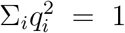. By defination the quaternion can be represented by 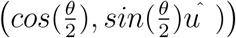, where *θ* is the optimum rotation angle and *û* is a unit vector associated with the optimum axis of rotation.

To estimate the distance between two orientation quaternion collective variables, (*q* − *q′*)^2^ = *g*^*ij*^(*q* − *q′*)_*i*_(*q* − *q′*)_*j*_, we use our approximation for the metric in (4.1). Following Dill (cite Dill’s paper), the orientation quaternions should satisfy the eigenvalue equation *Sq* = *λq*, where *S* is a 4 *×* 4 symmetric, traceless matrix, made of the correlation matrix (Euclidean coordinates). The *λ* are the eigenvalues *S*, and corresponding to eigenvectors (*q*). Differentiating both sides of the above eigenvalue equation we have:

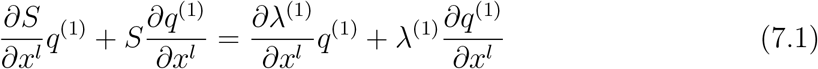

here *q*^(1)^ is the corresponding eigenvector to the leading eigenvalue *λ*^(1)^ = *λ*_*max*_. Using the orthogonality property between einvectors we obtain:

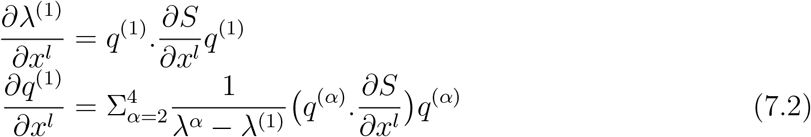

We can express the derivative of quaternions as

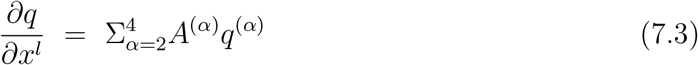

where we drop the index (1). So we can approximate our metric as

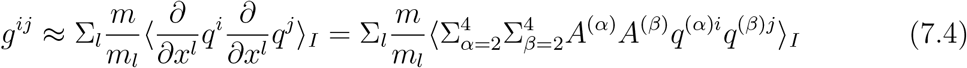

